# Transcriptional and cellular maturation of the chick spinal cord in the context of distinct neuromuscular circuits

**DOI:** 10.1101/2025.05.26.656077

**Authors:** Fabio Sacher, Bianka Berki, Antoine Fages, Libby Gavrilov, Avihu Klar, Maëva Luxey, Patrick Tschopp

## Abstract

The chicken spinal cord is a classic model system to study the early specification of neuronal cell types along its anterior-posterior axis. Here, we follow the ensuing maturation dynamics at limb levels with single cell resolution and contrast neuronal populations innervating appendages of distinct form and function. We use gene co-expression modules to identify rare cell populations with specific biological functions, and show that appendages with different motor outputs – wings and legs – rely on largely similar spinal cord cell type repertoires. Challenging the system with experimental alterations to the peripheral limb musculature reveals limited transcriptional changes, but spatially restricted plasticity in spinal cord motor neuron numbers. Collectively, our results provide a resource to investigate the molecular and cellular basis of neuronal maturation in the avian spinal cord and highlight the plastic nature of embryonic cells to adapt to changes in the limb periphery at both developmental and evolutionary timescales.

**Highlights:**

- Cellular and transcriptional maturation of the developing chicken spinal cord at single-cell resolution
- Single-cell weighted gene co-expression network analysis isolates transcriptional maturation dynamics in different neuronal populations
- *In silico* identification and *in situ* verification of novel markers for cerebrospinal fluid-contacting neurons
- Molecular characterization of the avian glycogen body
- Transcriptional and compositional changes of spinal cord cell types in the context of different neuromuscular circuits

## INTRODUCTION

The vertebrate spinal cord plays a critical role in coordinating motor control, relaying inputs from higher-order brain centers to the musculature, while receiving and pre-processing sensory feedback from the periphery. This bidirectional neuronal communication is crucial to ensure accurate movement and relies on the formation of precise neuromuscular circuits during development.^1–3^ At limb levels, motoneurons (MNs) located in the ventral horn of the spinal cord send out their axonal projections to connect to their corresponding muscles, to control voluntary movements of the appendages.^4,5^ Once initial contacts have been established, activity-based feedback from the periphery is critical to refine and mature these circuits, with sensory input communicated via the dorsal root ganglia (DRGs) to the mediolateral premotor interneurons.^6–8^ Interneurons and glial cells, including oligodendrocyte precursor cells (OPCs), are thus essential components in this maturation process. These cell types also contribute to the myelination of axons, to enhance the efficiency of signal transmission and support the long-term maintenance and functionality of neuromuscular circuits.^9,10^ Furthermore, other glial cell types such as astrocytes and microglia contribute to the maturation and the homeostasis of the system, by providing metabolic support and modulating its inflammatory responses.^11–13^ Accordingly, specification of a diverse set of neuronal and glial cell types, and their dynamic interplay with peripheral feedback mechanisms, is fundamental to the maturation of the spinal cord and its proper integration with the limb neuromuscular system.

The molecular programs establishing spinal cord cellular diversity are initiated early in development. Two sets of opposing morphogen gradients initiate neural tube patterning along both dorso-ventral and rostro-caudal axes, resulting in the establishment of distinct progenitor domains.^14–16^ Each functional territory is subsequently defined and subdivided by the combinatorial and highly specific expression of distinct sets of transcription factors (TFs), leading to the differentiation of the appropriate neuronal pools and cell types, with additional cellular diversifications occurring along a temporal axis.^17–19^ These early stages of spinal cord development have been well characterized in multiple species, and rely on a largely conserved molecular logic that instructs the specification of neuronal and non-neuronal cell types across vertebrates.^20–22^ Recent studies using single-cell RNA sequencing (scRNA-seq) have provided key insights into the genome-wide transcriptional dynamics underlying these early differentiation steps *in vivo* and *in vitro*.^20,23,24^ The ensuing maturation steps, however, and how they are affected by crosstalk with peripheral systems, remain largely unexplored.

The developing chicken spinal cord represents a particularly attractive model to study these later phases. Namely, the presence of appendages with distinct motor outputs – that is, wings and legs – offers an opportunity to study neuronal maturation in the context of neuromuscular circuits with distinct muscular topographies and neuronal firing patterns.^25,26^ Furthermore, we have the ability to induce limbs with extra digits and corresponding muscle bundles, thus providing us with an experimental setting to test for potential plasticity in response to an altered limb periphery during this maturation phase.^27,28^

Here, we investigate the transcriptional dynamics of chick spinal cord maturation at single cell resolution, from embryonic day 5 to day 10, and contrast neuronal populations innervating appendages of distinct motor outputs and muscle compositions. We demonstrate the power of gene co-expression modules to isolate and identify molecular signatures of rare cell populations with distinct biological functions, including the cerebrospinal fluid-contacting neurons (CSF-cNS) involved in controlling body posture and the avian-specific glycogen body. Furthermore, we show that chicken fore- and hindlimbs – i.e., wings and legs – rely on largely conserved cell type repertoires. Finally, by experimentally altering the skeletal formula and muscle patterns of the forelimb, we find limited transcriptional changes in neuronal cells, but spatially restricted alterations to MN cell numbers, highlighting the plastic nature of the limb-level nervous system to adapt to peripheral changes. Collectively, our data serve as a resource for future investigations into the transcriptional and cellular dynamics of spinal cord differentiation and maturation, and how these processes are affected by neuromuscular circuits of distinct form and function.

## RESULTS

### Single-cell profiling of the maturing chicken spinal cord across time and space

To follow the transcriptional and cellular dynamics of the developing chicken spinal cord during its maturation stages, we generated and sequenced 10x Chromium single-cell 3’ gene expression libraries obtained from micro-dissected tissue at brachial, forelimb-innervating levels, spanning three developmental stages from embryonic day 5 to day 10 (B05, B07, and B10, corresponding to Hamburger-Hamilton stages HH26, HH30, and HH36) (Figures S1A-C).^29^ Additionally, at embryonic day 10, we sampled and sequenced cells of the spinal cord at lumbar levels (L10), to investigate the cellular composition and transcriptional signatures associated with hindlimb-innervating neurons (Figure S1C). Finally, to probe for the potential of cellular and transcriptional plasticity in response to an experimentally altered peripheral limb musculature, we sampled and sequenced cells of the right half of brachial level-spinal cords that projected their axons towards a polydactyl wing (Poly10) (Figure S1D).^27^ For all samples, cells showed comparable distributions of unique molecular identifiers (UMIs), gene detection rates, and fraction of mitochondrial reads (Figures S1E-G). In total, we sequenced over 34,000 cells (2,474 – 6,866 cells per replicate), with an average of 3,293 genes detected per cell (346 – 6,894 genes). Using Louvain algorithms, we clustered our data sets individually and – based on differential gene expression (DE) analyses and known marker genes – broadly classified the resulting clusters according to the cell types they predominantly consisted of (Tables S1 and S2).^30^ Across our samples, pseudobulk transcriptome-based principal component analysis (PCA) first separated progenitor cells from maturing neurons, followed by the oligodendrocyte lineage and various non-neuronal cell types (Figures S1H and S1I).

We first focused our analyses on the temporally staggered samples at brachial levels. At 5 and 7 days of incubation, clusters belonging to either neural progenitor cells (NPC, yellow) or maturing neurons (magenta) were most abundant (Figures 1A and 1B), as evidenced by their *SOX9* and *TUBB3* expression (Figures 1D and 1E). Maturing neurons could further be sub-divided into excitatory or inhibitory types, based on the expression of *TLX3* and *NRXN3* (Figures 1D and 1E).^31,32^ Furthermore, we identified additional small, but distinct neuronal populations – such as roof plate (RP, green, *RSPO1*^+^) and floor plate (FP, maroon, *SHH*^+^) cells – as well as several non-neuronal populations such as pericytes (orange), blood (grey), and microglial cells (black) (Figures 1A-F). At embryonic day 10, oligodendrocyte precursor cells (OPC, blue) were more abundantly sampled, and one of the descendant lineages, the *PLP1*-expressing myelin-forming oligodendrocytes (MFOL, dark blue), was detected (Figures 1C and 1F).^33^ Along our developmental timeline we observed a depletion of neuronal progenitor fates (NPCs), in agreement with an ongoing cellular and transcriptional progression of spinal cord ontogeny, and the appearance of novel cell fates such as MFOLs, indicative of cellular maturation (Figure 1G). The relative scarcity of neurons in our day 10 sample could potentially be accredited to cytotoxic stress, resulting from axonal damage during dissection and tissue dissociation.^34^

**Figure 1.**
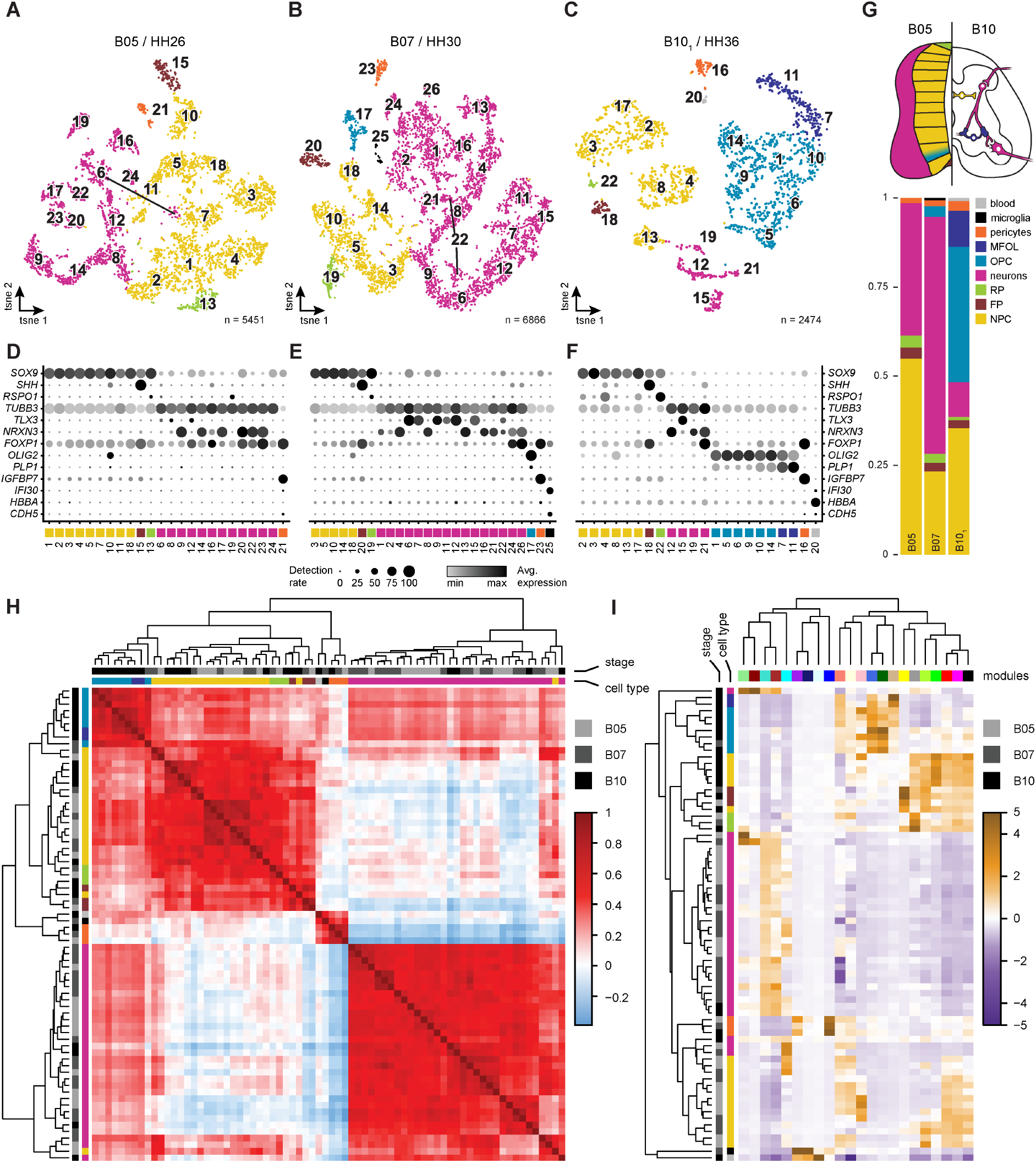
Transcriptional and cellular maturation of the brachial chick spinal cord. (**A-C**) tSNE embeddings of chick spinal chord samples from embryonic day 5 (**A**), day 7 (**B**), and day 10 (**C**). Colors indicate broad cell type categorization. (**D-F**) Dot plots showing marker gene expression across finer clustering, with dot size representing the fraction of cluster expressing a given marker, and gradient indicating the average scaled expression. Cell type colours correspond to Figure 1G. (**G**) Sketch shows spinal cord and approximate progenitor and neuron domains with bar plots of broad cell type contribution for all three stages. (**H**) Heatmap of spearman correlation of gene specificity indexed (GSI) pseudobulk expression. Log normalised gene expression from the SCT assay per time point is averaged by clusters and scaled by mean expression across sample (GSI). Only intersection of variable features identified by Seurat’s SCTransform() across all 3 time points (n = 1,841) were considered. Top row indicates time point, colour bar indicates broad cell type annotation. (**I**) Heatmap of average module eigengene expression by clusters and time point. Top row annotation indicates time point, colour bar indicates broad cell type annotation. Column annotation displays module color.

Despite these temporal differences in relative cellular abundances, the global transcriptional signatures appeared remarkably stable amongst functionally related cell types. Namely, we used gene specificity indices (GSI) of genes variably expressed across all samples, based on cluster-wise pseudobulk expression levels, to assess cell type-specific transcriptional similarities across embryonic timepoints.^35^ We calculated Spearman’s rank correlations of these GSIs and performed unsupervised hierarchical clustering. This revealed three major clades of transcriptionally related cells, regardless of the developmental stage that they were sampled from: progenitors and oligodendrocyte-related cells, non-neuronal cells, and maturing neurons (Figure 1H). This indicated that similar cell type repertoires were profiled along our sampling timeline. Accordingly, to further investigate the accompanying gene expression dynamics in these cells, we used single-cell weighted gene co-expression network analysis (scWGCNA) to construct modules of gene co-expression, and capitalized on their activity patterns to refine our cell type annotations.^36^ We integrated our single-cell gene expression data of samples B05, B07, and B10_1_ and identified a total of 22 gene co-expression modules that contained an average of 121 genes (16 – 485 genes) (Figure S2A, Table S3, and Table S4). Based on cellular module activity patterns, as well as associated enriched Gene Ontology (GO)-terms and KEGG pathways, we found that these modules appeared to represent a broad spectrum of cell type- and cell state-delineating gene activities, with varying degrees of specificity (Figures 1I, S2A, and S2B, Tables S5 and S6). For example, pericytes (orange) and microglia (black) populations were marked by the highly restricted activities of modules ‘blue’ and ‘midnightblue’, respectively (Figures 1I and S2B). Other modules’ activities outlined more general cell type classifications, such as modules ‘brown’ (GO:0048699, “generation of neurons”) and ‘turquoise’ (GO:0022008, “neurogenesis”) for developing neurons (magenta) (Figures 1I and S2B). Notably, the activity of additional modules allowed for a finer sub-classification of some of our more general cell type assignments. This is best illustrated by modules ‘darkred’ and ‘lightgreen’, which turned out to contain many genes known to be involved in motor neuron development, and thus allowed to identify these cells as a distinct neuronal sub-type (Figures 1I, 2A, 2D, and S2B). Additionally, some modules’ activities seemed to follow cellular maturation dynamics, even within a single sampling timepoint, as evidenced by modules ‘darkgreen’ and ‘tan’ that decreased and increased, respectively, as oligodendrocyte progenitors (OPCs) matured towards myelin-forming oligodendrocytes (MFOLs) (Figures 1I and S2B-D).

Collectively, our single-cell RNA sequencing sampling strategy of the chicken spinal cord, from early to mid-developmental timepoints, combined with unsupervised clustering, differential expression and scWGCNA gene co-expression analyses, demonstrated cell type conservation across our samples, and allowed for an evaluation of the cellular and transcriptional dynamics of cell type and cell state changes during spinal cord maturation.

### Cell fate- and cell state-defining gene co-expression dynamics during spinal cord maturation

Given the cluster-restricted activities of the scWGCNA modules ‘lightgreen’, ‘darkred’, ‘darkgreen’ and ‘tan’ (Figure S2B), we decided to investigate the associated molecular and cellular functions of their module members in more detail, as well as follow the temporal expression dynamics along the respective cell maturation trajectories.

Module ‘lightgreen’ contained the classical motor neuron markers *ALDH1A2, CHAT*, and *SLC18A3* (Figure 2A).^37–39^ The module also included *ISLR2*, a gene involved in axonal extensions in motor neurons, as well as *SEMA3C*, an axon guidance cue, with its receptor *NRP2* (Figure 2A).^40,41^ We removed redundant GO and KEGG pathway terms using REVIGO, to visualize a global semantic summary of the module activities.^42^ With respect to the other modules, the motor neuron module ‘lightgreen’ showed signs of enrichment in regulatory and stimulus-response terms, as well as in metabolism-related genes (Figure 2E). In contrast, the second motor neuron-specific module (‘darkred’) showed enrichment in homeostasis- and cellular organization-related terms (Figure 2F). This was also reflected in module gene composition, with acetylcholinesterase anchors *COLQ* and *PRIMA1*, as well as *SNCA* and *CPLX1* involved in presynaptic signaling (Figure 2B).^43–45^ Module ‘darkred’ thus reflected a more mature cell state of motor neurons than the one represented by module ‘lightgreen’. In agreement with this, the motor neuron cluster-specific expression levels of the two modules showed largely opposite activity patterns along our developmental timeline (Figures 2I and 2J).

**Figure 2.**
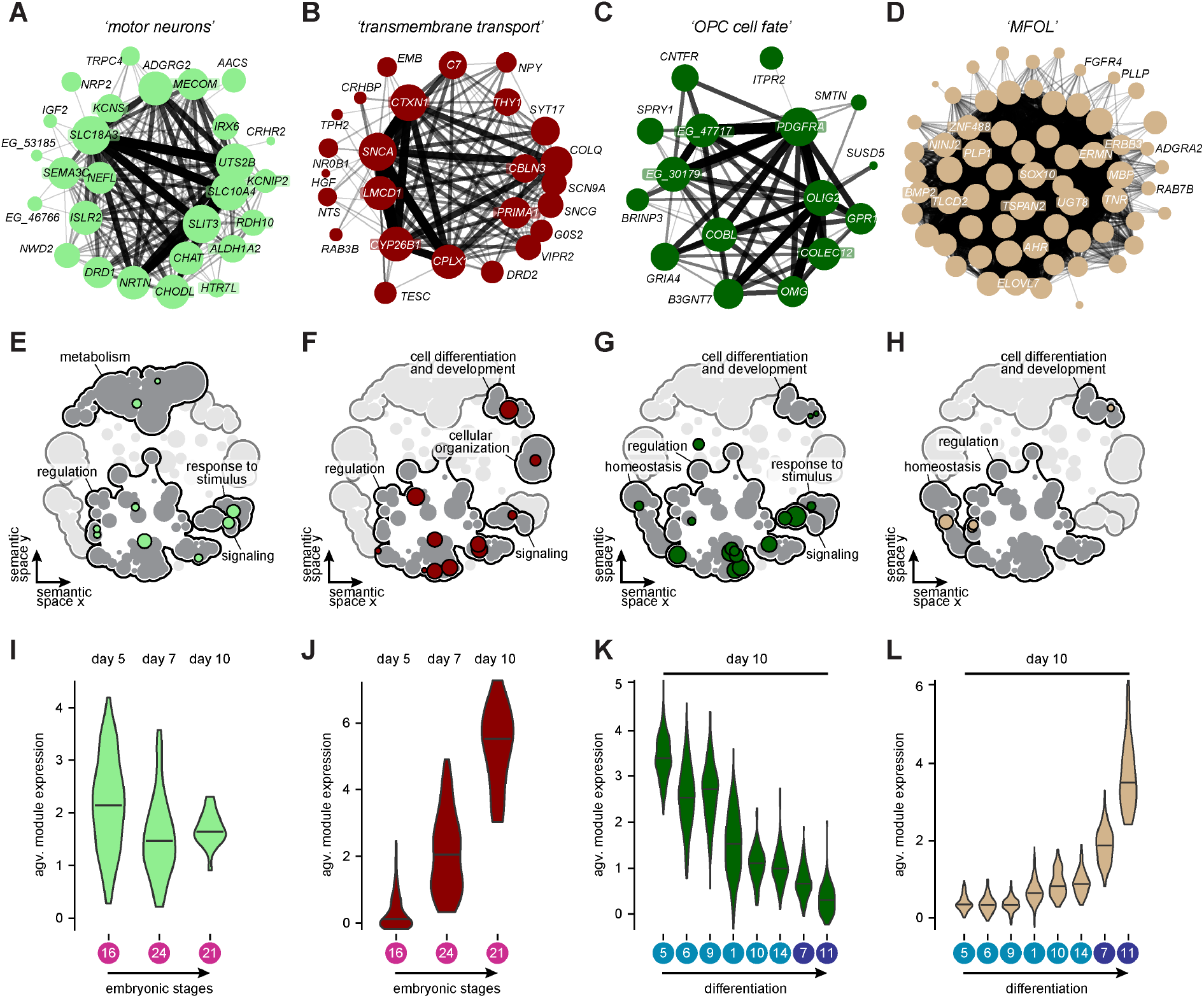
Gene co-expression dynamics of spinal cord cell type maturation. (**A-D**) scWGCNA module graph representations. Node size represent module membership and edge width and opacity represent topological overlap of expression. Uncharacterized genes labeled by “*EG_*”, followed by the last 5 digits of their *ENSGALG* gene codes. (**E-H**) Scatterplot of all modules’ GO Terms semantic space with module specific terms highlighted in module color. (**I-L**) Average module expression by broad cluster and stage. Black lines indicate median. (**I**,**J**) Average module expression of motor neuron related modules ‘lightgreen’ and ‘darkred’ per cell by motor neuron clusters across development. (**K**) Average module expression of the OPC related cluster ‘darkgreen’ per cell by OPC and MFOL cluster. (**L**) Average module expression of the MFOL related cluster ‘tan’ per cell by OPC and MFOL cluster. GO = Gene ontology, OPC = Oligodendrocyte progenitor cell, scWGCNA = single-cell Weighted gene co-expression network analysis.

The OPC clusters-labeling module ‘darkgreen’ was characterized by *OLIG2* and *PDGFRA*, two well-known oligodendrocyte precursor marker genes, as well as *OMG*, the oligodendrocyte myelin protein (Figure 2C).^46^ The module also included *SPRY1* and *SMTN*, two genes previously associated with spermatogonial stem cells and smooth muscle cells, respectively, that are transiently expressed in OPCs.^47,48^ Besides core regulatory terms, genes related to signaling and cellular differentiation were found to be enriched in this module (Figure 2G). The presence of *PDGFRA* indicated that the precursors marked by module ‘darkgreen’ expression had not yet differentiated into mature oligodendrocytes.^49^ In agreement with this, we found a decrease in module activity in clusters along the putative OPC-to-MFOL trajectory of our B10 sample (Figures 2K, S2C, and S2D). Conversely, module ‘tan’ showed an opposite activity pattern across the same oligodendrocyte clusters (Figures 2L, S2C, and S2D). The genes it contained – e.g. *MBP* (Myelin Basic Protein), *PLP1* (Proteolipid Protein 1), and *SOX10* – and their associated GO terms and KEGG pathways suggested a role in oligodendrocyte maturation, towards a cell state specialized in ramping up myelin-production (Figures 2D and 2H). In agreement with this, these putative MFOL clusters with increased levels of module ‘tan’ expression also displayed the highest activity of module ‘salmon’ (GO:0006412, “translation”), which relates to general protein synthesis (Figure S2B).^50,51^

Thus, using scWGCNA we were able to resolve distinct cell types and cell states, and follow gene expression dynamics along their respective differentiation trajectories, both within and across our sampling time points. Importantly, module activities seemed particularly suited for delineating sub-populations with different cellular and molecular functions, even when these cells types were only sparsely represented in our data.

### Identification of rare cell populations and associated marker modules

Capitalizing on the power of gene co-expression modules in identifying rare cell sub-populations, we performed iterative scWGCNA runs on our three replicated and integrated data sets at embryonic day 10: brachial (B10int), lumbar (L10int), and polydactyl (Poly10int) (see Figures S1C and S1D). This analysis resulted in 21, 50, and 19 modules, respectively (Figures S3A-C, Tables S3 and S4), to complement and refine our previous cell type annotations. Similar to our developmental time course, we found modules highly specific for non-neuronal cells – such as pericytes, blood, or vasculature – as well as modules delineating our broader neuronal classifications, like progenitors, maturing neurons, or oligodendrocytes (Figures S3A-C, Table S3). scWGCNA also identified modules labeling small but discrete cell populations, such as roof and floor plate cells (Figures S3A-C, black dashed rectangles), even though they accounted for less than 5 percent of the total cell numbers in each individual sample. These modules contained known markers of the respective cell populations, such as *LMX1A* for the roof plate or *SHH* for the floor plate, as well as cilia-related genes expressed in these same clusters (Figures S3A-D). Furthermore, in sample Poly10int, we identified a module ‘salmon’ whose activity appeared highly specific for a single neuronal cluster of only 63 cells (Figure S3C, arrowhead and salmon dashed rectangle). This module contained *PKD2L1*, as well as two pairs of paralogous transcription factors, *GATA2/3* and *TAL1/2* (Figure 3A). These genes had previously been identified in mice and zebrafish as markers for the so-called cerebrospinal fluid-contacting neurons (CSF-cNS), a population of specialized mechano- and chemosensory neurons that reside in the vicinity of the central canal and project their ciliated apical extensions into the cerebrospinal fluid.^52–55^

**Figure 3.**
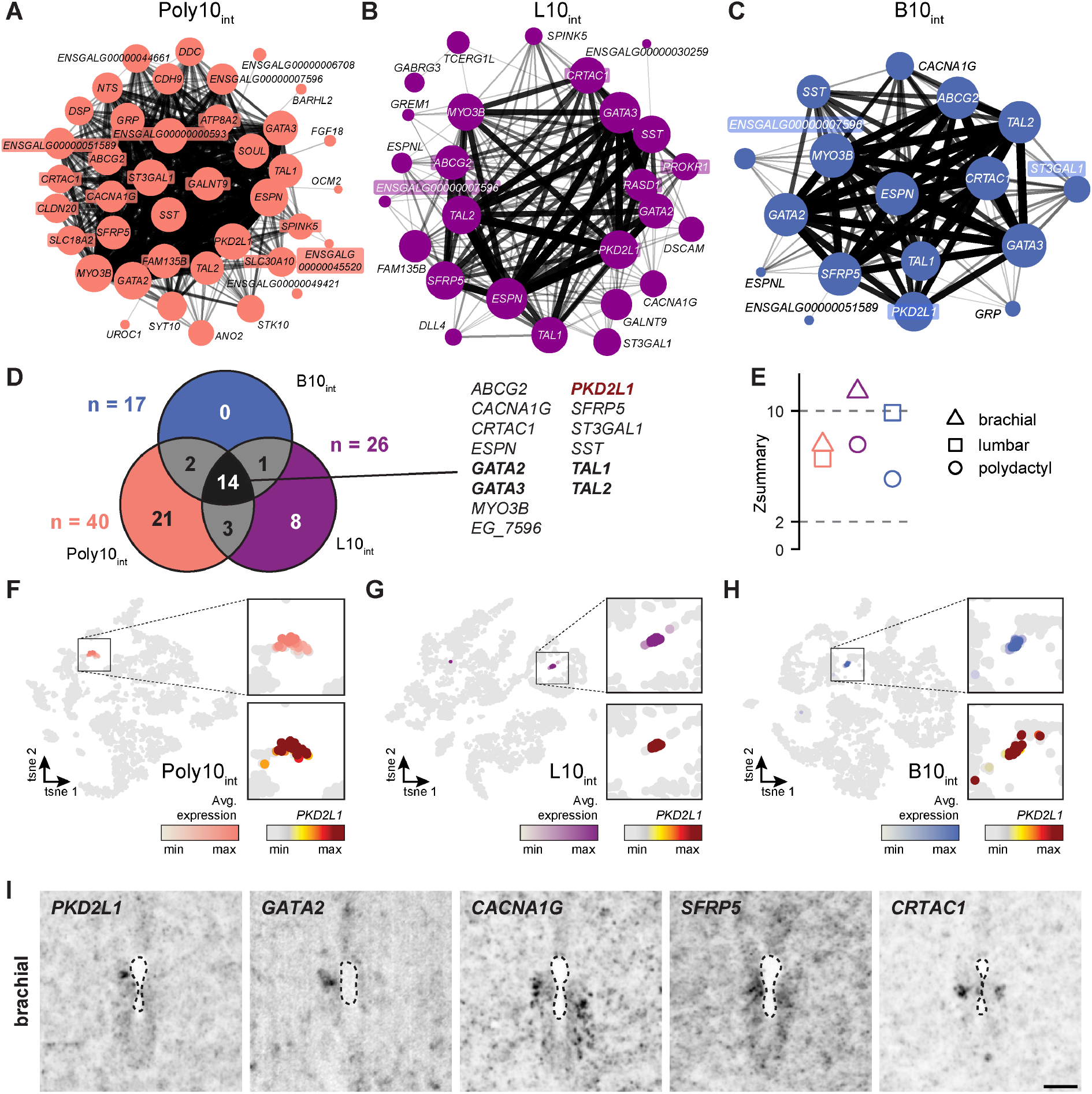
Transcriptional signatures of the cerebrospinal fluid-contacting neurons. (**A-C**) scWGCNA module graph representations of the CSF-cNS related modules in the polydactyl (**A**), lumbar (**B**), and brachial (**C**) samples. Node size represent module membership and edge width and opacity represent topological overlap of expression. Uncharacterized genes labeled by “*EG_*”, followed by the last 5 digits of their *ENSGALG* gene codes. (**D**) Venn diagram showing module size and intersection of all three modules. Known transcription factors involved in CSF-cNS development in italics. (**E**) scWGCNA Z-summary of module conservation across samples. Thresholds indicate weak (2) and strong (10) indication for module conservation. (**F-H**) tSNE embeddings showing average module expression by cell and scaled marker expression of *PKD2L1*. (**I**) *In situ* hybridization of CSF-cNS associated genes (*PKD2L1* and *GATA2*) and novel candidates (*CACNA1G, SFRP5*, and *CRTAC1*) on brachial day 10 spinal cord sections. Images show inverted fluorescent channel. Min and max values adjusted by crop. Scale bar, 50 μm. CSF-cNS = cerebrospinal fluid-contacting neurons.

We identified two additional modules, ‘royalblue’ in B10int and ‘darkmagenta’ in L10int, whose gene contents showed substantial overlap with module ‘salmon’ (Figures 3B-D), with elevated expression levels in one neuronal cluster each of the respective samples (Figures S3A and S3B, black arrowheads and royalblue and darkmagenta dashed rectangles). These three modules shared a total of 14 genes, including somatostatin (*SST*), expressed in lamprey lateral CSF-cNS, as well as *ESPN* and *MYO3B*, an actin bundling gene and its activator known to be expressed in zebrafish CSF-cNS (Figure 3D).^56–58^ Hence, using scWGCNA, we identified a core module set of markers for the CSF-cNS spinal cord population, containing known and novel candidate genes of diverse molecular functions (Figure 3D), with cluster-enriched expression patterns (Figures S3A-C) and preserved module ‘densities’ and ‘connectivities’ (Figure 3E).^36,59^

Module activities, as indicated by the module eigengene, colocalized nicely with *PKD2L1* expression (Figures 3F-H), as well as with newly identified marker candidates (Figure S3E, arrowheads). To evaluate potential CSF-cNS-related expression patterns *in vivo*, we performed fluorescent *in situ* hybridization on cryosections of day 10 brachial-level spinal cords for *PKD2L1* and *GATA2* transcripts, as positive controls, and *CACNA1G, SFRP5*, and *CRTAC1*, as novel putative marker genes. High levels of gene expression always were detected adjacent to the central canal, where CSF-cNS population resides, thus confirming our *in silico* predictions *in situ* (Figures 3F-I and S3E).

Collectively, through combinatorial scWGCNA analyses, we identified modules of co-expressed markers for distinct neuronal and non-neuronal populations, with varying degrees of cell type- and cell state-specific molecular activities. Importantly, we were able to isolate and transcriptionally characterize a rare cell population of cerebrospinal fluid-contacting neurons (CSF-cNS), which would not have been identified using conventional clustering and DE methods.

### Cell type repertoires in spinal cord segments innervating wings and legs

We next explored potential cellular and molecular differences in spinal cord segments innervating wings and legs. Chicken appendages are used for drastically different modes of locomotion. While hindlimbs for walking and running show the alternating gait mode present in most terrestrial tetrapods, the coordinated bilateral flapping of their forelimbs propels powered flight. The required synchronization of the underlying contraction patterns in birds has been linked to distinct circuit architectures at brachial levels and the natural loss of function of an axon guidance molecule.^26,60^ However, whether differences in these neuromuscular circuits are also reflected at the cellular or molecular level has not been investigated in detail yet. Accordingly, to probe for putative cellular and molecular differences in spinal cord segments mediating either flying or walking motor behaviors, we compared our day 10 brachial (wing-innervating) and lumbar (leg-innervating) spinal cord data sets (Figure S1C).

We first integrated and annotated our duplicate brachial and lumbar samples separately, which resulted in 23 and 29 clusters, respectively (Figures S4A and S4B). We recovered similar cell type repertoires at the two anatomical locations, as evidenced by marker gene expression indicative of the major classifications (Figures S4C and S4D) and sample-wise pseudobulk PCA (Figures S1H and S1I). Overall, we observed a trend for a higher fraction of neuronal progenitors and reduced numbers of maturing neurons, OPCs and MFOLs at lumbar levels, reflecting the general heterochrony of the spinal cord along its anterior-posterior axis (Figure S4E).^61^

To mitigate potential technical sampling effects, and to directly compare cells between the two spinal cord locations, we integrated all brachial and lumbar samples together and re-clustered them, revealing a total of 34 clusters (Figure 4A). Similar to B10int (Figure S4A), RP and FP cells formed a single cluster (B/L10int-12), potentially driven by the shared expression of the cilia/flagella modules (Figure S3D). Across all integrated clusters we only found minor transcriptional differences between brachial and lumbar cells, even though some differentially expressed axon guidance molecules might be instructive for generating distinct wiring patterns at fore- and hindlimb levels (Figure S4K, Table S7). Using the R package miloR, we then tested for differential abundances of brachial versus lumbar cells in our integrated B/L10int data (Figure 4A).^62^ This revealed four clusters showing strongly imbalanced sample representations in their miloR neighborhoods (Figure 4B, asterisks). First, the motor neuron-containing cluster B/L10int-30 consisted predominantly of brachial cells, as expected from our failure to detect a distinct cluster for this cell type in our lumbar samples (Figures 4B, S4A (B10int-20) and S4B). Second, cluster B/L10int-3 displayed a bimodal distribution of miloR abundance changes, with different parts of the cluster being dominated by either brachial or lumbar cells (Figures 4A and 4B). Elevated expression of *PBX3, MEIS1*, and *SLC17A6* indicated that this differentially abundant sub-cluster likely consisted of differentiating glutamatergic cells of the dorsal horn (Figure S4F).^63^ Lastly, clusters 21 and 22 showed the strongest abundance biases, towards the lumbar samples (Figure 4B). Shared gene expression signatures – of transcripts differentially expressed in these two clusters versus the rest – indicated a molecular affinity with roof and floor plate cells (Figure 4C). Besides genes involved in cellular host defense, such as *GSTA4* or *FIBCD1*, the expression levels of the branching enzyme *GBE1*, Enolase-1 (*ENO1*), an enzyme involved in glycolysis, and *PPP1R3B*, a phosphatase subunit regulating glycogen synthesis, were also elevated in B/L10int clusters 21 and 22 at lumbar levels (Figure 4C).^64^ This suggested that these cells might belong to the glycogen body, a glial structure in the lumbar spinal cord of birds.^65,66^ This was confirmed by *in situ* hybridizations on lumbar cryosections, using probes against *GBE1* and *MSX2*, two transcripts enriched in clusters 21 and 22, which showed high expression levels in the developing glycogen body (Figures 4C and 4D). Additionally, two scWGCNA modules identified in our lumbar samples showed their highest expression in these two clusters. While ‘magenta’ appeared restricted to glycogen body-related clusters B/L10int-21 and -22 (Figures 4E and S3B, red arrowheads and dashed rectangle), ‘brown4’ displayed a broader activity pattern, in neurons as well as the glycogen body and roof plate cells (Figures S4G and S3B, dashed rectangles). This allowed us to investigate the transcriptional signatures of developing glycogen body cells in greater detail. The smaller module ‘brown4’ (n = 13 genes) contained *CRMP1* – a nervous system-specific cytosolic phosphoprotein involved in semaphorin-signaling – and the transcription factor *ZIC1* (Figure S4H).^67^ The much larger module ‘magenta’ (n = 73 genes) included additional markers associated with a dorsal or roof plate fate, such as *PAX3* and MSX1, as well genes connected to various neurological and neurodegenerative disorders like *FAM222A* (e.g. Alzheimer’s disease) and *NDP* (e.g. retinopathies) (Figure S4I).^68,69^ Both modules also contained multiple members of the family of *HOX* gene transcription factors (Figures S4H and S4I). Due to the axial differences between the brachial and lumbar spinal cord segments, we had expected strongly divergent *HOX* gene expression signatures.^70^ Indeed, most neurons and oligodendrocytes showed the expected bimodal distribution of anterior and posterior *HOX* genes expression, between our samples of brachial and lumbar origins (Figure S4J). This was particularly evident for the *HOXD* cluster, whose expression – besides *HOXD3* and *HOXD4* – was largely absent from neuronal cells at brachial levels and oligodendrocytes throughout (Figure S4J, red dotted frames).^71,72^ Strikingly, however, the four module members *HOXD3, HOXD4, HOXD9* and *HOXD10*, as well as their genomic neighbor *HOXD8*, were all expressed at lumbar levels, and particularly enriched in our glycogen body cells, as was *HOXA11* of module ‘magenta’ (Figures 4F and S4J, green shades). This suggested that the lumbar glycogen body cells carry a distinct *HOX* code, with a particular enrichment for members of the *HOXD* cluster.

**Figure 4.**
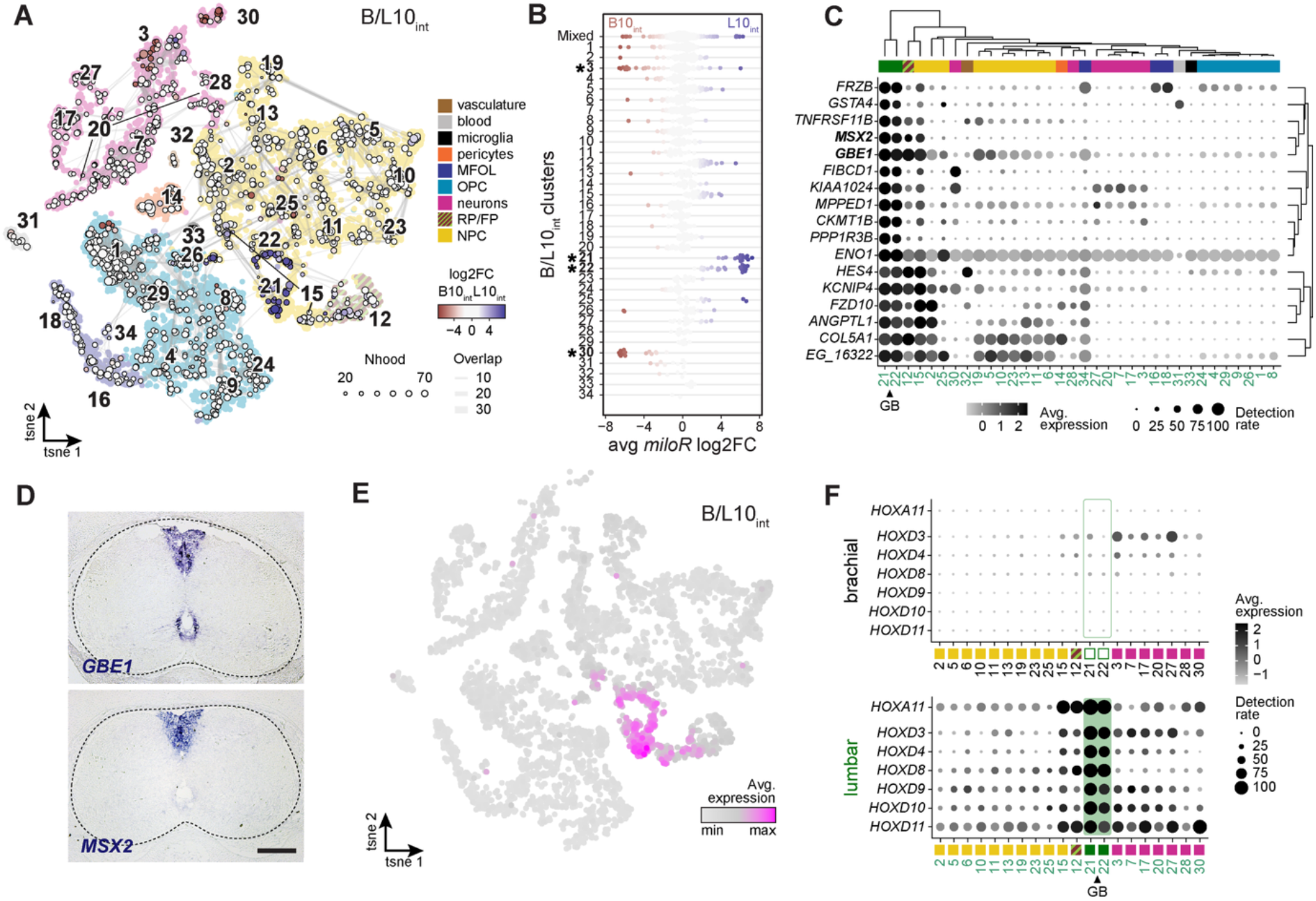
Cell type repertoires at wing- and leg-innervating spinal cord segments. (**A**) tSNE embeddings of integrated embryonic day 10 brachial and lumbar (B/L10_int_) chick spinal cord samples with network of differentially abundant miloR-neighborhoods superimposed. Base colors indicate broad cell type categorizations. FP/RP cluster contains cells from both populations. (**B**) Beeswarm plot of differentially abundant neighborhoods by cluster. Asterisks indicate the most biased clusters. (**C**) Dot plot of intersected top differentially expressed genes for glycogen body (GB) clusters 21 and 22. GB clusters are indicated in dark green. Uncharacterized genes labeled by “*EG_*”, followed by the last 5 digits of their *ENSGALG* gene codes. (**D**) *In situ* hybridization of *GBE1* and *MSX2* on day 11 lumbar chick spinal cord, showing expression in the forming glycogen body. Scale bar, 200 μm. (**E**) tSNE embedding showing average expression of module ‘magenta’. (**F**) Dot plots of selected *HOX* gene expression for progenitors, neurons, FP, RP, and GB. The plots are split by brachial and lumbar. log2FC = average logarithmic fold change FP = floor plate, RP = roof plate.

Collectively, our comparative analyses of the brachial and lumbar spinal cord revealed largely conserved cell type repertoires, with corresponding cell populations from the anterior and posterior regions showing only minor transcriptional changes. A notable exception was the identification of cells forming the glycogen body, an avian-specific structure of the lumbar region that showed a distinct enrichment for *HOXD* transcripts.

### Plastic responses in motor neuron numbers to an experimentally altered limb periphery

Besides intrinsic differences in spinal cord wiring patterns, brachial and lumbar segments also differ in the axonal projections of their motor neurons, which eventually connect to appendage-specific muscular topographies in the fore- and hindlimb peripheries.^26,27^ Given that – on an evolutionary timescale – muscle and innervation patterns needed to change in a coordinated manner, to ensure continuous functionality of the full limb neuromuscular system, this suggests a certain degree of developmental plasticity in generating the underlying circuits, to accommodate putative changes in one system by the other.^28,73^ We have previously documented such plasticity at the morphological level, in both peripheral muscle and innervation patterns, using different polydactyly models in the chicken.^27,28^ Here, we capitalized on this system to probe for cellular and molecular plasticity in spinal cord segments projecting to an experimentally altered limb.

Our samples were obtained from bisected brachial spinal cords, split into control (B10) and experimental polydactyl (P10) sides (Figures 5A and S1D). Our integrated polydactyl samples (P10int) revealed 29 clusters, with similar cell type repertoires and proportions as for our control counterparts (Figures S5A-C). A notable exception was the over-representation of floor and roof plate cells, likely resulting from our midline bisection (Figure S5A). To quantify possible differential cell abundances across our experimental conditions, we integrated duplicate control and polydactyl samples and ran miloR. As expected, both roof plate (B/Poly10int-22) and floor plate (B/Poly10int-23) clusters had many neighborhoods with a pronounced overrepresentation of polydactyl-side cells (Figures 5A and 5B). Furthermore, the closely related NPC neighborhoods in clusters 16 to 19 all showed slight polydactyl enrichments, although these imbalances were again likely due to the inclusion of roof or floor plate-like cells, as a result of our bisection approach (Figures 5A and 5B). Overall, compositional cell number biases were also apparent in the number of differentially expressed genes we detected between control and polydactyl sides, with strong abundance differences being mirrored by more differentially expressed genes (e.g. clusters B/Poly10int-18 and 22, Figures 5B and 5C, asterisks, Figure S5D, Table S7). Clusters with similar abundances across samples (miloR |log2FC|< 1) only showed minor transcriptional changes, as exemplified by the neuron clusters B/Poly10int-5 (inhibitory) and 11 (excitatory), or the motor neurons in B/Poly10int-25 (Figure 5B, asterisks, Figure S5D).

**Figure 5.**
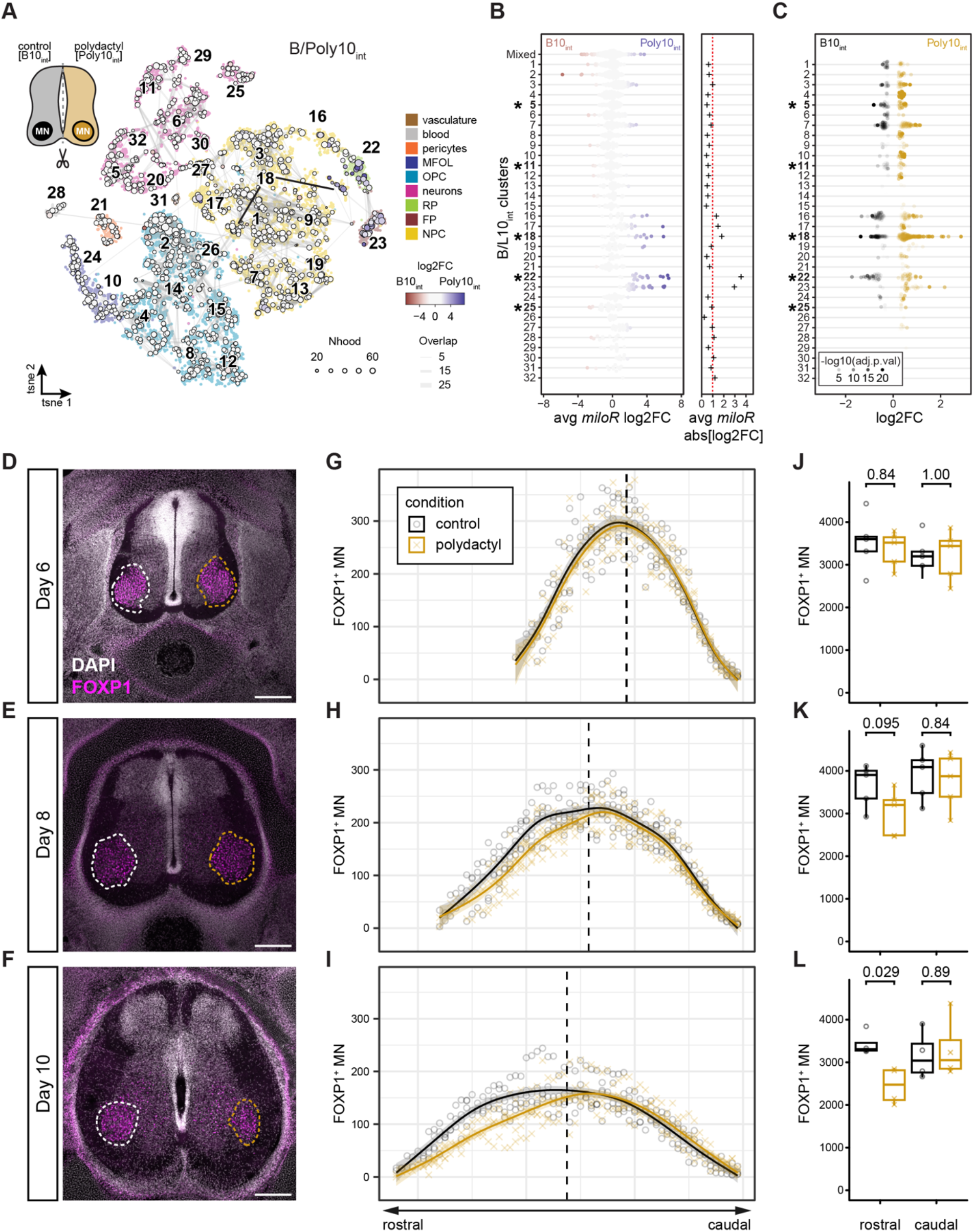
Spinal cord cellular plasticity in response to an altered limb periphery. (**A**) tSNE embedding of embryonic day 10 integrated brachial and polydactyl (B/Poly10_int_) spinal cord samples with network of differentially abundant miloR-neighborhoods superimposed. Base colors indicate broad cell type categorization. Schematic of bisection procedure on top left. (**B**) Beeswarm plot of differentially abundant neighborhoods by cluster, with absolute miloR log-fold changes shown on the right. Asterisks indicate clusters with high (cl-18 & cl-22) or low (cl-5 & cl-11) abundance changes, and motor neurons (cl-25). (**C**) Beeswarm plot of differentially expressed genes between brachial and lumbar cells by cluster. Alpha values represent −log10 of adjusted p-values. (**D-F**) Representative sections per day stained for FOXP1 and DAPI. Dashed lines indicate LMCs. Scale bar, 200 μm. (**G-I**) Counts of FOXP1+ nuclei per section (50μm) at day 6 (n = 5), 8 (n=5), and 10 (n=4) between control and polydactyl brachial LMC. Dashed line indicates split between rostral and caudal halves. (**J-L**) Boxplots of average motor neuron count split by day, condition, and axis. Brackets indicate compared groups and p-values as calculated by Wilcoxon test.

We next investigated potential effects of changes in limb musculature on motor neuron cell fates in a more targeted fashion. Given that proper wiring is essential for neuromuscular circuit function, motor neurons that fail to establish proper connections to their target muscles are selectively culled over the course of development.^74^ Accordingly, we assessed spinal cord motor neuron numbers separately for either control or polydactyl sides, by quantifying FOXP1-positive nuclei on cryosections at embryonic day 6, 8, and 10 (Figures 5D-F).^75^ Furthermore, we capitalized on their limb segment-specific rostral-caudal arrangement along the lateral motor column (LMC), whereby motor neurons towards the rostral end of the brachial LMC are known to innervate more proximal target muscles, while digit-innervating cells are located at its caudal extremity.^76^ Across the three embryonic stages, using 4-5 experimental replicates each, we quantified over 191,000 motor neurons along the rostral-caudal axis of the brachial LMC (Figures 5G-I). At day 6 of development, motor neuron numbers on control and polydactyl sides of the same spinal cord looked indistinguishable (Figure 5G). Over the following four days, however, we observed a clear decline of motor neuron numbers in the rostral halves of the polydactyl sides (Figures 5H and 5I). These rostral changes became more pronounced over time, whereas – unexpectedly – no significant changes were observed in the caudal, digit-innervating end (Figures 5J-L).

Overall, despite the experimental induction of substantial changes in the peripheral limb musculature, we uncovered only minor transcriptional differences in limb-level spinal cord cells. However, we found a clear reduction of rostral LMC motor neurons upon the polydactyl-induced alteration of muscle topography, indicating that periphery-related spinal cord plasticity at this stage manifests itself predominately at the level of cell numbers.

## DISCUSSION

The vertebrate nervous system has long served as a prime model system to study cell fate diversification across both developmental and evolutionary timescales. By capitalizing on the power of single-cell functional genomics methods, it is now possible to track the molecular dynamics of its ontogenetic specification with unprecedented resolution, as cells undergo complex and multilayered maturation processes, from initial stem cell fate decisions to plastic neuronal adaptations in cell states. Once the major distinct neuronal cell type classes have been specified, spinal cord neurons involved in the control of voluntary movement become embedded in neuronal circuits that play an increasingly important role in their ensuing maturation. Here, using droplet-based single-cell transcriptomics, we follow the molecular and cellular maturation dynamics of chicken spinal cord neurons, from late cell type specification stages to early circuit refinements, in the context of distinct peripheral neuromuscular circuits.

### Delineating cell types and cell states during spinal cord maturation

By combining unsupervised clustering analyses with known cell type marker gene expression profiles and *de novo* gene co-expression modules reconstruction, we catalogued the cellular diversity of the maturing chicken spinal cord at single-cell resolution. Along our developmental sampling timeline at brachial levels, we documented a progressive decrease in uncommitted neuronal progenitors, with a concomitant increase of maturing neurons and glial cells. We detailed the ongoing refinement of distinct cell types – such as the progressively restricted delineation of FOXP1+ motor neurons (Figures 1A-F) – as well as the late appearance of entirely new and distinct cellular functions, such as the onset of myelin-ensheathment by oligodendrocytes (Figure 1G). We demonstrate the power of reconstructing scWGCNA gene co-expression modules in demarcating different clades of related cell types, some of which with partially overlapping sets of module activities such as translation, while more restricted ones allowed for a finer scale of cell sub-type or state classifications (Figure 1I). The module-associated GO terms and KEGG pathways enable quick and cluster-specific insights into the presumptive biological, cellular and molecular players at work – such as the involvement of the cilia machinery in floor and roof plate cells (Figures S3A-D) or their spatial location (Figure S2B, ‘grey60’, GO:0048263 “dorsal identity” and ‘yellow’, GO:0009950 “dorsal/ventral axis”).^77^ Furthermore, they can help to define novel candidate genes involved in orchestrating the ongoing cell type and cell state diversification during spinal cord maturation (Figure S2B, Table S4). Importantly, averaged module activities can reveal the underlying developmental dynamics of these maturation processes, both across as well as within sampling timepoints (Figures 2 and S2B-D). For example, while early cell type specification modules ‘lightgreen’ and ‘darkgreen’ decreased over developmental time, in motor neurons and oligodendrocytes, respectively, modules ‘darkred’ – enriched for cation transmembrane transport and neuroactive ligand-receptor interaction – and ‘tan’ – neuron insulation and glial maturation – increased in activity, indicative of a cell state maturation specific to their underlying cell types.^78^ Building on such well-studied cell types as a proof-of-principle allowed us to identify additional co-expressed genes and associated molecular functions for them – e.g. “translation” (GO:0006412, module ‘salmon’) in myelinating oligodendrocytes (Figure S2B) – or to investigate module activities in rare cell types lacking high-resolution molecular data so far.

### Annotating rare cell types using gene co-expression modules

Using scWGCNA we identified several gene co-expression modules highly specific for cell populations of low relative abundance, such as e.g. pericytes (module ‘blue’) or microglia (modules ‘midnightblue’ and ‘purple’) (Figure S2B). Furthermore, using an iterative approach, we identified several modules with overlapping gene compositions and transcriptional activities that delineated the cerebrospinal fluid-contacting neurons (CSF-cNS) (Figures 3A-H).

CSF-cNS are specialized type of neurons, located close to the central canal of the spinal cord, which probe the cerebrospinal fluid via their ciliary projections. By mechano- and chemosensory modalities, they act as cellular detectors that respond to fluctuations in the state of the cerebrospinal fluid, as well as the spine curvature itself.^55^ For example, CSF-cNS are able to detect pathogen metabolites and show an increase in Ca2^+^ activity upon CSF infection.^79^ Furthermore, via their interaction with the Reissner fiber, they sense changes in the spatial conformation of the spinal cord and are critical for maintaining body posture during zebrafish development.^80,81^ While their various physiological functions are thus increasingly appreciated, here we present extensive gene co-expression module data that identify novel markers for these elusive cells.^17,82^ For instance, we detected module co-expression of *ESPN* and *MYO3B* – hearing- and deafness-related genes identified in mice and zebrafish CSF-cNS – thus indicating a conserved molecular machinery in cilia-based mechano-sensation between CFS-cNS and inner ear hair cells.^83,84^ Finally, using fluorescent *in situ* hybridization, we verify CSF-cNS-specific module expression adjacent to the central canal using known and novel marker genes (Figure 3I).

Not only did scWGCNA help us to isolate and characterize rare cell populations, it also enabled the molecular description of a phylogenetically restricted cell type. Cells of the glycogen body – a dorsal, glycogen-rich glial structure that arises from the roof plate cells, present at sciatic levels of the lumbar spinal cord in birds and possibly dinosaurs – showed enrichment for various enzymes involved in sugar metabolism (Figures 4C and 4D).^26,65,85^ Their confined rostral-caudal localization at sciatic lumbar levels, together with the neuronal circuits that mediate left/right limb motor control, support the hypothesis that the glycogen body evolved to enable alternating stepping movements in avian hindlimbs.^26^ Additionally, scWGCNA identified two gene co-expression modules enriched in glycogen body cells that contained multiple transcriptional regulators, including a highly specific set of *HOX* gene transcription factors (Figures 4F and S4H-J). These distinct regulatory codes can inform us about the underlying transcriptional changes that define the clade-specific phylogenetic occurrence of these cells, as well elucidate their ontogenetic relationship with the adjacent roof plate. Thus, future experimental manipulations of glycogen body-specific transcription factors may help to reveal the role of the glycogen body in the uncoupling of synchronized wing flapping and legs alternation, which is a unique characteristic of avians.

### Plastic cellular responses to peripheral changes in muscle topography

Despite the distinct locomotory modes and muscle patterns of the appendages they innervate – i.e., wings and legs – the overall cellular composition and their transcriptional signatures in brachial and lumbar spinal cord segments appeared remarkably conserved (Figures 4A, 4B, S4C, and S4D). This supports a serially homologous nature of tetrapod appendages also at the level of spinal cord cell type repertoires, upon which avian clade-specific modifications were implemented.^26,65,66,86^ Furthermore, it suggests that evolutionary changes in motor firing patterns are primarily driven at the circuit level, while relying on similar cell types.^26,60,87,88^ Likewise, we only detected minor transcriptional changes in spinal cord cells connecting to either control or polydactyl wings (Figures 5C and S5D). This lack of plastic molecular differentiation was surprising, given that our experimental intervention is known to introduce substantial alterations to the limb musculature, and the important role of muscle-derived peripheral feedback on neuronal maturation.^27,89–91^ The low number of surviving motor neurons in our data set, due to potential dissociation stress, might have prevented us from detecting more subtle and motor neuron pool-specific transcriptional changes. Relying on single nuclei-based approaches, to improve motor neuron survival, or using motor neuron-targeted techniques, with much higher gene detection sensitivity, may help to reduce these issues in future studies.^92,93^

We did, however, identify spatially restricted changes in motor neuron numbers, indicative of a segment-specific response in the central nervous system to peripheral changes in muscle topography. Such cellular plasticity in the tetrapod spinal cord might have facilitated the evolutionary coordination with adaptive changes in limb morphologies, as distinct muscular and skeletal formulas evolved in response to locomotory specializations.^94,95^ Globally, the decrease in motor neurons largely followed previously described proportions, with slightly lower overall cell numbers being distributed over an expanding spatial domain (Figures 5G-I and S5E).^96,97^ Importantly, however, motor neuron numbers were significantly and specifically lowered at the rostral ends of our polydactyl-side LMCs, compared to control side numbers (Figures 5I and 5L). This apparent reduction can potentially be explained by a loss of proximal-anterior muscles – replaced by posterior duplicates – and, therefore, a lack of innervation targets for the rostrally located motor neurons that should normally innervate these limb areas.^27,89,98^. Yet, despite the presence of multiple additional distal autopod muscles in our polydactyl wings, we only detected minor, non-significant increases in caudal motor neuron numbers on the polydactyl sides (Figure 5I and 5L).^27,76^

## Conclusions

Collectively, we provide a cellularly resolved transcriptional resource to follow the maturation dynamics of the chick spinal cord. We delineate the molecular changes that underlie cell type specification and cell state changes, using gene co-expression modules for broad and rare cell populations. Furthermore, by contrasting spinal cord segments innervating appendages with distinct neuromuscular circuits and motor outputs, we define conserved and plastic cellular responses to peripheral changes relevant at developmental and evolutionary timescales.

## MATERIAL & METHODS

### Induced polydactyly in chicken

Fertilized chicken eggs (Gallus gallus) were acquired from vendors in Switzerland. Eggs were incubated at 38°C with 60% humidity for 3 days, then windowed and staged according to the Hamburger-Hamilton developmental table.^29^ Resin beads (AG1-X2, Biorad) were formatted in formic acid for an hour, and washed with clean water until the pH stabilized around 4.5. Before implantation, beads were incubated in 1 mg/ml all-trans retinoic acid (RA) solution for at least 20 min, then incubated in DMEM media (Gibco) until they turned red. A small incision was made between the Apical Ectodermal Ridge and the mesenchyme at the anterior margin of the limb bud and RA-soaked beads were implanted.^27^ Eggs were closed with a tape and incubated until the desired stage (3, 5, or 7 days after operation). In this study, only individuals with digit patterns 432234 were included.

### Sample dissociation and single-cell RNA sequencing preparation

Untreated embryos were harvested at embryonic day 5 (HH27, B05), 7 (HH30, B07), and 10 (HH36, L10_1_, L10_2_) and Polydactyl embryos were harvested at day 10 of development (HH36, B10_1_, B10_2_, Poly10_1_, Poly10_2_) in ice cold PBS, decapitated and transferred into a black bottomed SYLGARD™ Petri dish. They were pinned facing down, an incision was made in the middle of the back and dorsal laminectomy was performed to get access to the neural tube. For the polydactyl embryos, roof plate and the floor plate were cut with a needle, before dissecting the part corresponding to the caudal LMC. Control and polydactyl side were separated into different tubes and processed separately. Dissected caudal LMCs were transferred into 3 ml of 37°C warm dissociation media (FACSmax™ media, Amsibio with 0,375 mg/ml Papain, Millipore Sigma) and incubated for 5 min in a shaking dry bath at 37°C and 300 rpm. The first three triturations were performed with a P1000 pipette tip cut 7, 5, and 3 mm from the tip and quickly burnt with a Bunsen burner to eliminate sharp edges. The last trituration was carried out with an uncut P1000 pipette tip. Each trituration was followed by 5 min of incubation at 37°C. Cell suspensions were centrifugated at 300 g for 7 min at 4°C. The supernatant was discarded, cell pellets were resuspended in 1 ml HBSS 5% FBS (ThermoFisher), passed through a 40 μm cell strainer (Flowmi®, Millipore Sigma) into a 1,5 ml Eppendorf tube and centrifuged again at 300 g for 7 min at 4°C. Cell pellets were resuspended in 100 μl HBSS 5% FBS and cell number and viability were assessed with an automated cell counter (Cellometer®, AOPI ViaStain™). Only samples above 80% viability were processed for sequencing. Samples were loaded on Chromium Next GEM Chip G following the manufacturer’s instructions (Chromium Next GEM Single-cell 3 Reagent Kits v3.1 (Dual Index); available on https://support.10xgenomics.com), aiming for the recovery of 10,000 cells. cDNA and libraries were prepared following the above-mentioned protocol, cDNA and library qualities were assessed with a Bioanalyzer (Agilent). The sequencing was carried out in the D-BSSE (ETH Zürich, Basel) at the Genomics Facility.

### QC and filtering

The raw sequencing data was processed with Cell Ranger software (10X Genomics) version 3.1.0 (B10_1_, B10_2_, L10_1_, L10_2_, Poly10_1_, Poly10_2_) and version 5.0.1 (B05, B07) to perform base calling, adapter trimming, and mapping to the GRCg6a chicken genome assembly and its extended annotation, published by Feregrino & Tschopp.^36^ Cell filtering steps were applied depending on sample quality and sequencing depth, applying thresholds excluding cells with more than 4 times the mean UMI counts or less that 20% of the mean, as well as removing cells with a mitochondrial UMI fraction bigger than the median + three times the MAD (median absolute derivation). Cells with a gene to UMI ratio smaller than 15% and less expressed genes than 2/3 of the maximum number of detected genes were also not considered for further analysis. In total, we obtained 34,340 high quality transcriptomes (2,474 – 6,866 cells per sample).

### Seurat objects

With Seurat v4.0.5, cells passing the thresholds were imported as Seurat objects by sample.^99^ With SCTransform() the objects were then normalized and scaled, regressing out mitochondrial percentage (mt.perc) and number of UMI counts. Cell cycle stages G2M and S were scored with Seurat’s CellCycleScoring() to calculate their difference (CC.difference), that was then regressed out together with the previous variables and the sample ID in another run of SCTransform prior to principal component analysis (PCA). The chicken cell cycle markers were obtained by selecting orthologous genes from the provided human genes. Principal component dimensions for t-SNE were selected for each sample as those lying outside of a Marchenko-Pastur distribution (histogram of squared standard deviations) and used in fftRtsne() (Fast Fourier transform accelerated interpolation based t-SNE) to calculate a two dimendional embedding.^100,101^ Cell clusters were identified with FindNeighbours() and Find-Clusters(), using the same dimensions identified above. Based on a tree of clusters in PCA space, terminal sister clusters with less than 20 differentially expressed highly variable genes (var > median(var)) between them are merged. Marker genes are obtained using FindAllMarkers() with the parameters: only.pos = TRUE, min.pct = 0.25, logfc.threshold =0.5, latent.vars = c(“CC.difference”), test.use = “MAST”, assay = “RNA”, and return.thresh= 0.05. The features used are the var.features of the SCT assay after running SCTransform again regressing out mt.perc, UMI counts, and sample ID.

### Dot plots

Dot plots were created with a modified version of Seurat’s DotPlot(), available from the R package modplots on GitHub (https://github.com/safabio/modplots).

### Integration

The Devel_int_ (B05, B07, B10_1_), B10_int_ (B10_1_, B10_2_), L10_int_ (L10_1_, L10_2_), Poly10_int_ (Poly10_1_, Poly10_2_), B/L10_int_ (B10_1_, B10_2_, L10_1_, L10_2_), B/Poly10_int_ (B10_1_, B10_2_, Poly10_1_, Poly10_2_) and All_int_ (all samples) objects were integrated following the SCTransform integration vignette. Then, cell cycle scoring, scaling with ScaleData() and vars.to.regress set to “CC.difference”, PCA, tSNE, and clustering was performed on the integration assay, following DE analysis on the RNA assay using the parameters described above.

### PCA of all samples

A RNA counts pseudobulk matrix of individual clusters across all samples in the All_int_ was obtained with Seurat’s AggregateExpression(). Using DESeq2, a DDS object was created with DESeqDataSetFromMatrix() and DESeq() and transformed with vst().^102^ PCA on the top 1000 variable genes was calculated and plotted with a modified version of DESeq2’s plotPCA, available from the R package modplots on GitHub (https://github.com/safabio/modplots).

### GSI Pseudo bulk correlation

To determine cell type relationships in Devel_int_, pseudo bulk of the SCT data slot was calculated with AverageExpression(). Average expression by cluster was divided by average expression across all clusters to obtain gene specificity indices. Heatmaps and hierarchical clustering of pairwise spearman correlation were created with pheatmap v1.0.12 (Figure 1E).^103^ Only the intersection of SCT variable features of the individual data sets were considered.

### scWGCNA

Modules of co-expressed genes for Devel_int_ (B05, B07, B10_1_), B/L10_int_ (B10_1_, B10_2_, L10_1_, L10_2_), and B/Poly10_int_ were constructed using the scWGCNA R package v1.0.0.^36^ The default assay was set to “SCT” and PCA was performed. Variable features from the integrated assay were set as SCT variable features before constructing modules following the package’s suggestions. GOterm enrichment analysis and KEGG pathways were performed with goana() and kegga() from the limma package v3.48.0.^104^

The Devel_int_ modules constructed on the integrated time series were used to calculate module eigengenes for each cell in the individual Seurat objects. The module eigengenes were averaged by cluster. The heatmap of the pseudo bulk module eigengenes was created pheatmap (Figure 1F). Violin plots were created with VlnPlot(), using average module expression as features and grouping by cell cluster (Figures 2I-L). To plot module expression by cell, the WGCNA data was split by sample and the averageExpr values from the sc.MEList slit was added to the meta data of each Seurat object. Then, tSNE embeddings highlighting the average module expression were obtained with FeaturePlot() (Figures S2B and S2C). Networks plots were created with adapted code from scWGCNA::scW.p.network (Figures 2A-D, 3A-C, S4H, and S4I). Module expression along development was plotted by splitting the WGCNA data by sample and plot the mean average expression values by broad cluster (Figure S2B). Comparison of modules across B10_int_, L10_int_, and Poly10_int_ done with scWGNA.compare() (Figure 3E). Module expression plots were done with scW.p.expression() (Figures 3F and 3G).

### Revigo

Semantic analysis of GOterm enrichment in Devel_int_ modules was conducted on http://revigo.irb.hr/ (Figures 2E-H).^42^ Revigo was run with parameter Tiny(0.4) against the GalGal database. GO terms were filtered to p.val < 0.01 and N terms was set to < 3000.

### Differential abundance

Differential abundance (DA) was calculated following the basic example of the miloR v1.0.0 workflow (Figures 4A, 4B, 5A, and 5B).^62^

### *In situ* hybridization

Control embryos were dissected at day 10 and fixed in 4% PFA overnight, cryoprotected in 30% Sucrose solution, embedded in (FSC22 Clear Frozen Section Compound, Biosystems) blocks and kept at −80°C until sectioning. 18 μm thick sections were cut on a Leica cryostat from caudal to rostral. Sections were air dried for an hour after and frozen at −20°C until *in situ* hybridization was performed using standard protocols.^105^ For fluorescent probe detection, TNT (0.1 M Tris pH 7.5, 0.15 M NaCl, 0.05% Tween) washes and peroxidase inactivation with 2% H_2_O_2_/TNT, was followed by blocking with 0.5% BR/TNT (Akoya Biosciences) and incubation with 1:300 anti-DIG POD (Roche) in 0.5% BR/TNT. TSA Plus Cyanine 3 and TSA Cyanine 5 (Akoya Biosciences) was used for signal amplification, while NBT/BCIP was used for chromogenic revelation. Primers for RNA probe production are listed in Table S8.

Slides were imaged with a Leica DM4B fluorescent microscope using a 5X objective (N.A. 0.12, air, N Plan). The images were rotated so that the central canal was centered and vertical, cropped and adjusted with resetMinAndMax() in FIJI.

### Immunohistochemistry

Polydactyl embryos were dissected at day 6, day 8, and day 10, DiI was injected into the right side of the back to identify polydactyl sides later on sections. Embryos were fixed in 4% PFA overnight, cryoprotected in 30% Sucrose solution, then embedded in (FSC22 Clear Frozen Section Compound, Biosystems) blocks and kept at −80°C until sectioning. 50 μm thick sections were cut from caudal to rostral in order to capture the full length of the brachial LMC. Sections were air dried for an hour after and frozen at −20°C until immunostaining. Slides were thawed for 15 min, washed in PBS to eliminate OCT and blocked in PBST (0,1% Triton, 0.02% SDS, 0.002% BSA in PBS) solution for an hour. Primary antibodies against FOXP1 (Abcam) were diluted 1:500 in PBST and slides were incubated with the mix in a humid chamber overnight at 4°C. After three PBST washes, anti-rabbit AF647 secondary antibody (1:500) were deposited on the slides, incubated for 2 h at room-temperature, washed three times in PBST and mounted with fluorescent mounting media (Agilent). Slides were kept at 4°C until imaging.

### Confocal Imaging

Slides were imaged with an Olympus Fluoview (FV3000) confocal microscope using a 10x objective (N.A. 0.4, air, ApoPlan, Olympus). For FOXP1-positive cell counting, Z-stacks were acquired with 3 μm steps between virtual sections.

## ACKNOWLEDGMENTS

The authors wish to thank the “Genomics Facility Basel” for help with library preparation and sequencing, and all members of our group for useful discussions. All calculations were performed at sciCORE (**http://scicore.unibas.ch/**), scientific computing center at the University of Basel. This work was supported by funds from the Janggen-Pöhn Foundation to F.S., the Israel Binational Science Foundation (grant no. 1787/21) to A.K., and the Swiss National Science Foundation (SNSF project grant 310030_170022), the Olga Mayenfisch-Foundation, the FSRMM (Fondation Suisse de Recherche sur les Maladies Musculaires), and the University of Basel to P.T.

## AUTHOR CONTRIBUTIONS

F.S., B.B., M.L. and P.T. conceived the study. F.S., B.B., A.F., and M.L. performed experiments, F.S., B.B., M.L. and P.T. analyzed the data, L.G. and A.K. provided reagents and data, F.S., B.B., A.K., M.L. and P.T. interpreted data, F.S. and P.T. wrote the manuscript, with feedback from all co-authors.

## Supplementary Figures

**Figure S1.**
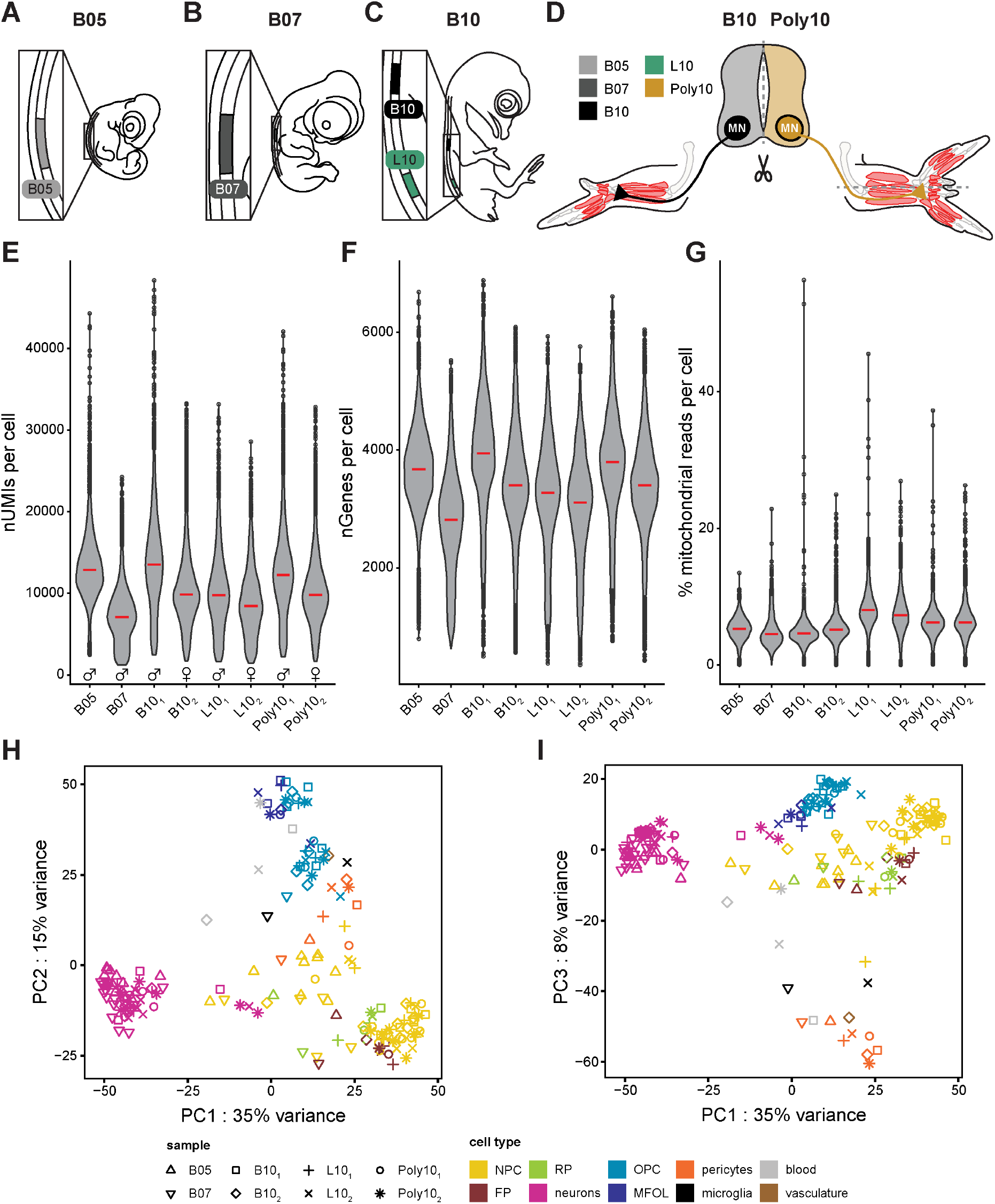
Samples collection and single-cell transcriptome quality control metrics. (**A-C**) Sampling strategy for day 5 (**A**), day 7 (**B**), and day 10 (**C**). (**D**) Sampling strategy for day 10 control and polydactyl spinal cords. (**E-G**) Violin plots of number of UMIs (**E**), number of detected Genes (**F**), and percentage of mitochondrial reads (**G**) per cells, split by individual sample data sets. Red lines indicate median. Sex of the individual embryos is indicated in E. (**H**,**I**) Principal component analysis of pseudobulk aggregate expression across all integrated samples. Cluster identities were transferred from the individual data sets. Shapes indicate samples. (**H**) Principal components 1 and 2. (**I**) Principal components 1 and 3.

**Figure S2.**
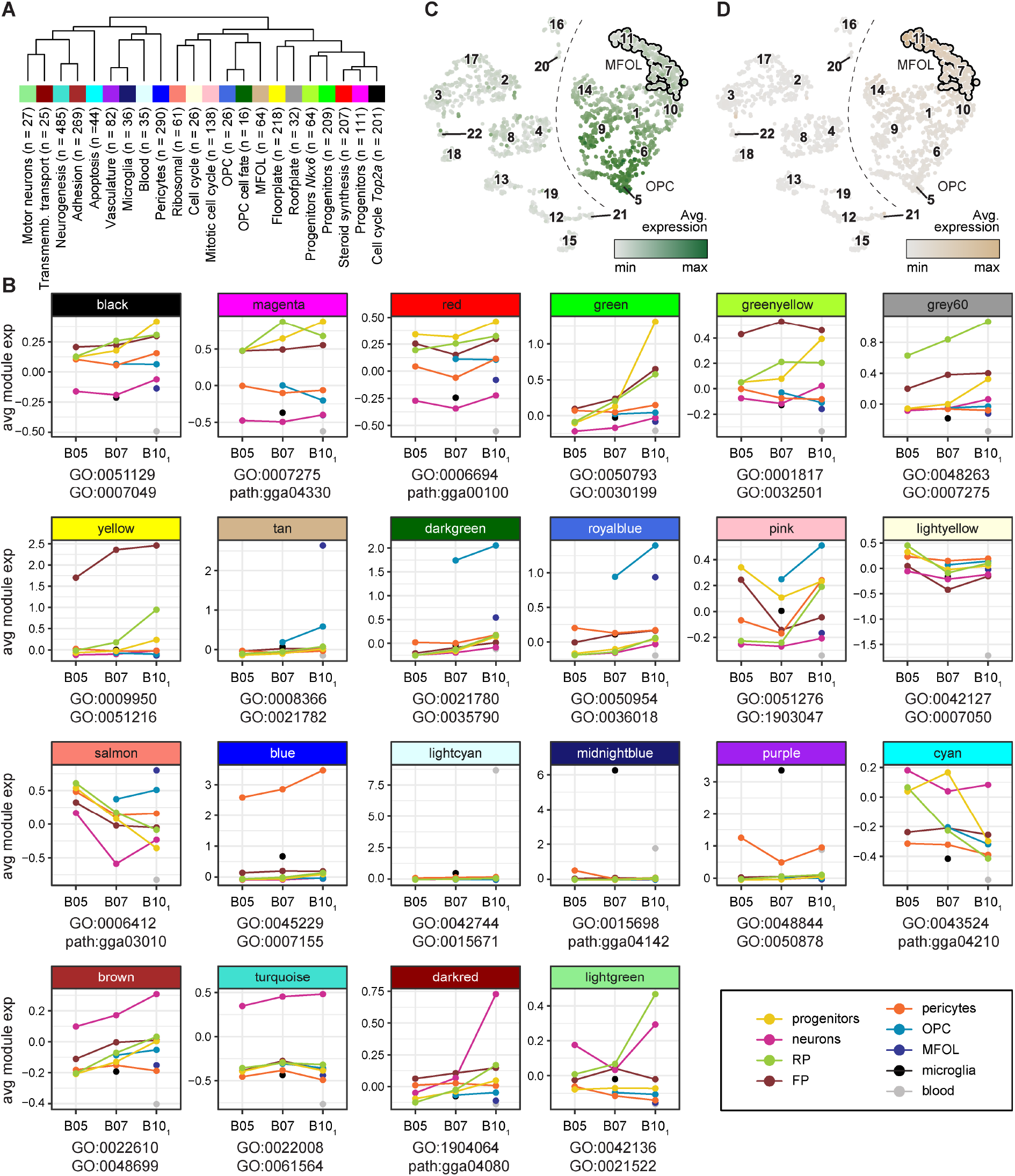
Gene co-expression modules of brachial spinal cord cell maturation. (**A**) Module tree of the average module eigengene expression heatmap shown in Figure 1F with module annotations and sizes indicated below. (**B**) Average module expression by broad cluster and stage. Relevant GO terms or KEGG pathways indicated below each plot. (**C**) tSNE representation of B10_1_. Color gradient shows average module expression of the OPC related module ‘dark green’. Dashed lines separate Oligodendrocytes from the remaining cell types. MFOLs are highlighted by a black line. (**D**) tSNE representation of B10_1_. Color gradient shows average module expression of the MFOL related module ‘tan’. Dashed lines separate Oligodendrocytes from the remaining cell types. MFOLs are highlighted by a black line. GO = Gene Ontology, MFOL = myelin forming oligodendrocytes, OPC = oligodendrocyte progenitor cell.

**Figure S3.**
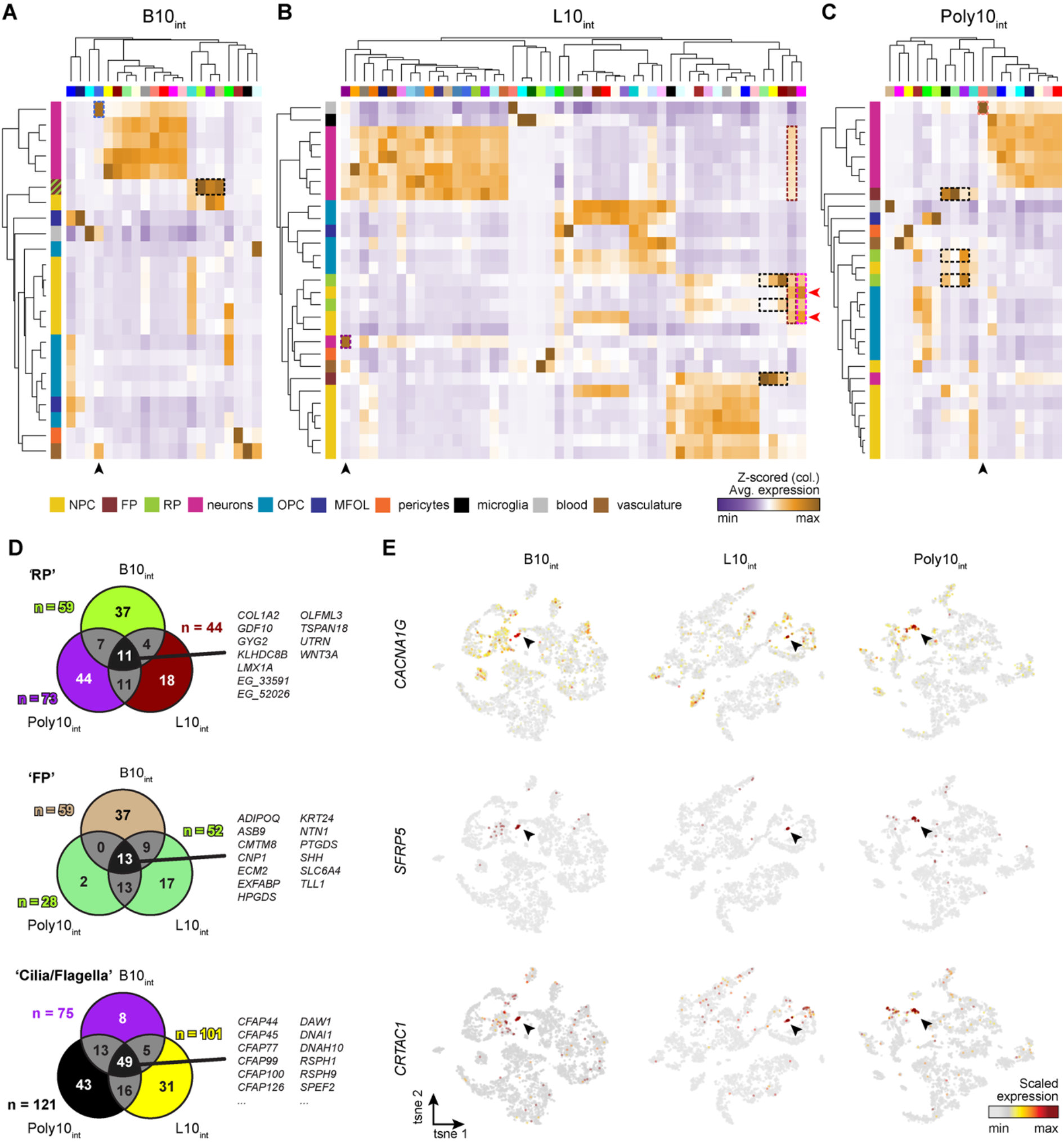
Gene co-expression modules of embryonic day 10 spinal cord samples. (**A-C**) Heatmap of average module eigengene expression by clusters. Black arrowheads indicate CSF-cNS modules. (**A**) B10_int_ modules. Dashed rectangles highlight expression of modules ‘royalblue’ (CSF-cNS), ‘greenyellow’ (RP), ‘tan’ (FP), and ‘purple’ (clilia/flagella). (**B**) L10_int_ modules. Dashed rectangles highlight expression of modules ‘darkmagenta’ (CSF-cNS), ‘darkred’ (RP), ‘lightgreen’ (FP), and ‘yellow’ (clilia/flagella). Red arrowheads indicate GB clusters (**C**) Poly10_int_ modules. Dashed rectangles highlight expression of modules ‘salmon’ (CSF-cNS), ‘purple’ (RP), ‘lightgreen’ (FP), and ‘black’ (clilia/flagella). (**D**) Venn diagrams showing module size and intersection of RP, FP, and clilia/flagella modules for B10_int_, L10_int_, and Poly10_int_. Uncharacterized genes labeled by “*EG_*”, followed by the last 5 digits of their *ENSGALG* gene codes. (**E**) tSNE embeddings showing scaled gene expression for the CSF-cNS markers *CACNA1G, SFRP5*, and *CRTAC1*. Arrowheads indicate CSF-cNS cells as identified by module expression in Figures 3F-H. FP = floor plate, RP = roof plate.

**Figure S4.**
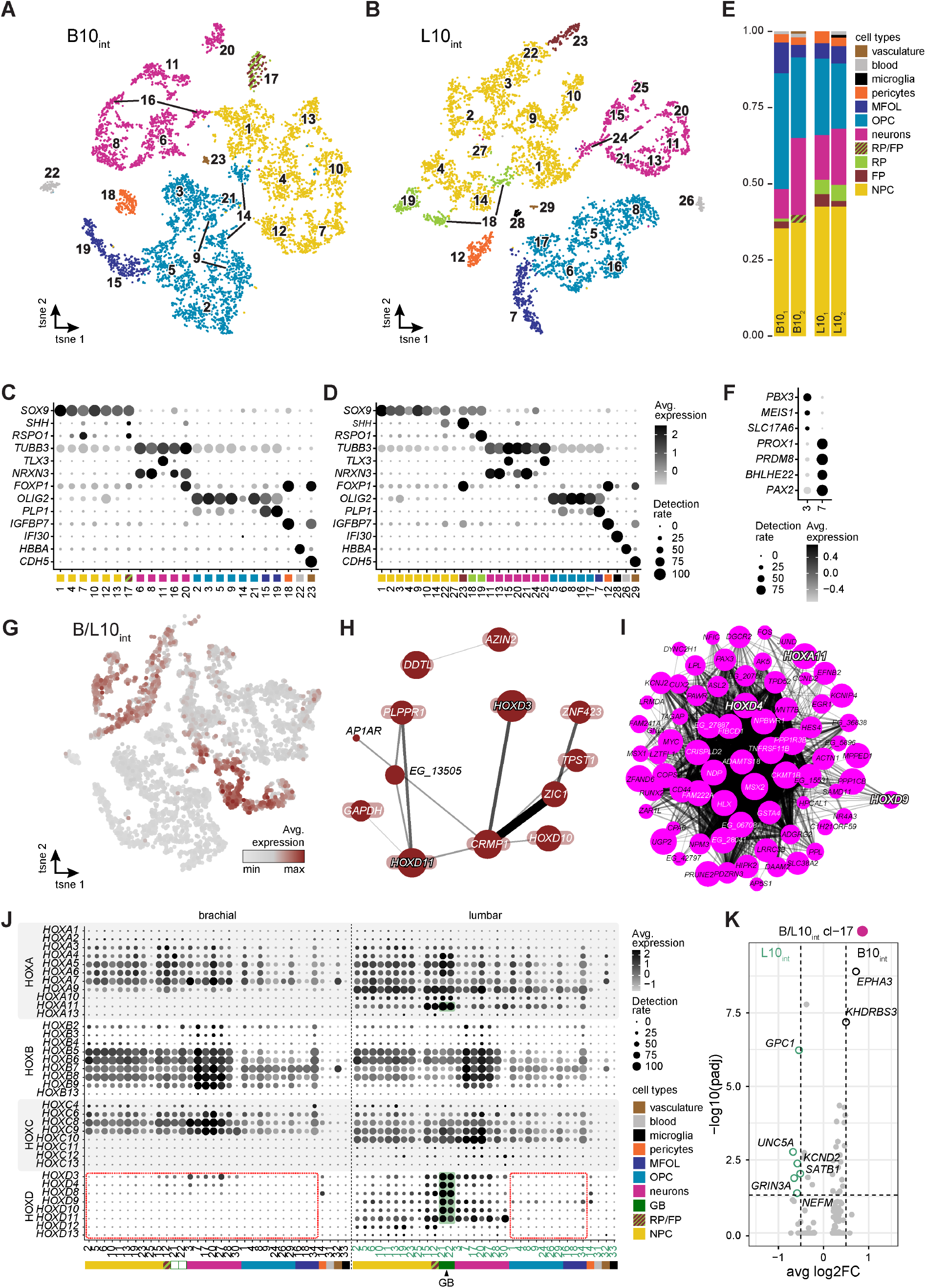
Cell type repertoires and gene expression signatures at wing- and leg-innervating spinal cord segments. (**A**,**B**) tSNE embeddings of brachial (**A**) and lumbar (**B**) chick spinal cord samples from day 10. Colors indicate broad cell type categorization (**C**,**D**) Dot plots show marker expression across finer clustering for day 10 brachial and lumbar integrated data sets. Marker selection was based on cluster annotation relevance. Dot size represents fraction of cluster expressing a given marker, gradient indicates average scaled expression. (**E**) Bar plot of broad cell type contribution for all 4 samples. (**F**) Dot plots of genes delineating the differentially abundant sub-cluster within B/L10_int_ cl-3 and from cl-7. (**G**) tSNE embedding showing the average expression of module ‘brown4’. (**H**,**I**) scWGCNA module graph representation of module ‘brown4’ (**H**) and ‘magenta’ (**I**). Node size represent module membership and edge width and opacity represent topological overlap of expression. *HOX* genes in white bold italic with black outlines. Uncharacterized genes labeled by “*EG_*”, followed by the last 5 digits of their *ENSGALG* gene codes. (**J**) Dot plot of *HOX* cluster genes split by brachial and lumbar cells. Red rectangle highlights absence of *HOXD* cluster genes in all brachial clusters and lumbar oligodendrocyte clusters. (**K**) Volcano plot of differentially expressed genes between B10 and L10 cells in B/L10_int_ cl-17.

**Figure S5.**
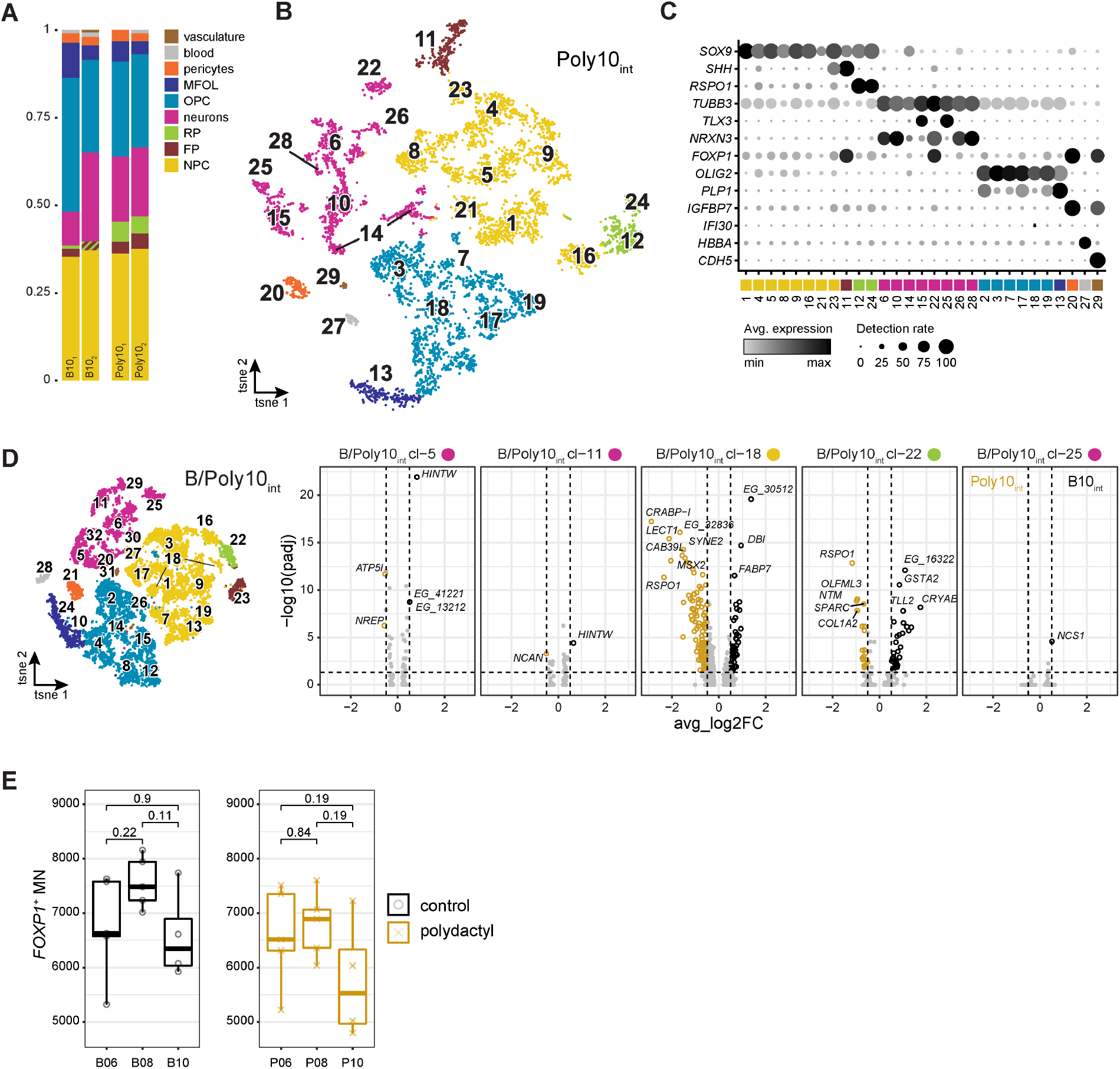
Spinal cord transcriptional and cellular plasticity in response to an altered limb periphery. (**A**) Bar plot of broad cell type contribution for all 4 samples. (**B**) tSNE embeddings of polydactyl chick spinal cord samples from day 10. Colors indicate broad cell type categorization. (**C**) Dot plots of marker expression across fine clustering of the integrated day 10 polydactyl samples. Color indicates broad cell clusters (**D**) tSNE embeddings of integrated brachial control and polydactyl chick spinal cord samples from day 10. Colors indicate broad cell type categorization. Volcanoplots of intra-cluster DE genes of the clusters highlighted with asterisks in Figure 5C. Uncharacterized genes labeled by “*EG_*”, followed by the last 5 digits of their *ENSGALG* gene codes. (**E**) Boxplots showing the total count of MNs per embryo and day for control and polydactyl sides. Brackets indicate comparisons with p-values calculated by the Wilcoxon test.

